# IgG4 serum levels are not elevated in cases of Post-COVID syndrome

**DOI:** 10.1101/2023.03.01.530454

**Authors:** Jonas Abel, Annika J. Walter, Vivian Glück, Clara L. Magnus, Thomas Glück, Philipp Schuster, Stefan Blaas, Ida Montanari, Michael Koller, Arno Mohr, Thilo Hinterberger, Bernd Salzberger, Kerstin Renner, Matthias Mack, Robert Bals, Tina Schmidt, Verena Klemis, Martina Sester, Romina Kardashi, Katja de With, Thomas H. Loew, Maximilian Malfertheiner, Michael Pfeifer, André Gessner, Barbara Schmidt, Daniel Schmalenberger, David Peterhoff, the COVIDYS consortium

## Abstract

Recently, unexpectedly high virus-specific IgG4 levels were reported after more than two mRNA vaccinations. Class switch towards IgG4 occurs after long-term antigen exposure, downregulates immune responses and is associated with several autoimmune diseases.

Here, we examined differences in antigen-specific IgG subtypes in serum samples from 64 Post-COVID patients and an equally sized cohort of convalescent controls.

In both cohorts, the relative amounts of spike protein-specific IgG subtypes were comparable. IgG1 was the most frequent, followed by IgG3, IgG2, and IgG4. A difference between cohorts was observed only for IgG2, which was significantly lower in the Post-COVID cohort. Further analysis of the reactive IgG4 revealed a small but significant difference for the spike protein receptor-binding domain but not for the spike ectodomain.

Since the total IgG4 levels are very low, we do not expect a biologically relevant role in Post-COVID syndrome. However, reduced virus-specific IgG2 levels could contribute to the persistence of SARS-CoV-2, causing chronic inflammation in the setting of Post-COVID syndrome.

---

Numerous studies have recently described levels and durability of infection-or vaccination-induced SARS-CoV-2-specific serum antibodies as a correlate of protection. In addition to their virus-neutralizing effect through receptor competition and stabilization of the viral spike fusion machinery, the importance of their effector functions is currently debated (1). Effector functions are closely linked to the four antibody subtypes (IgG1, -2, -3, -4), i.e. sequence, structure and glycosylation of the antibodies’ fragment crystallizable region (Fc) (2).

In this context, two independent publications recently described the finding of significantly increased IgG4 levels after more than two mRNA vaccinations, which was not seen in subjects vaccinated with adenoviral vectors (3,4). Antibodies of the IgG4 subtype occur after long-term antigen exposure, downregulating immune responses and inducing tolerance (2). Furthermore, IgG4 antibodies are associated with several autoimmune diseases (5). The authors suggested that mRNA-vaccine induced elevated IgG4-levels may be due to a prolonged availability of antigen in the germinal centers of the lymph nodes after mRNA vaccination or to a specific property of the mRNA vaccine. It is currently unknown whether IgG4 is also induced after SARS-CoV-2 infection in non-vaccinated individuals. Furthermore, the pathogenic potential of IgG4 in the context of SARS-CoV-2 vaccination and COVID-19 is unclear. This prompted us to investigate whether increased IgG4 levels also occur in patients suffering from Post-COVID syndrome. In these patients, prolonged circulation of viral antigen has been described (6). Increased class-switch of B-cells towards IgG4 might result in less Fc-mediated effector function and thus contribute to longer viral persistence and prolonged antigen circulation.

We examined this hypothesis in cohorts of 64 Post-COVID patients (20 males, 44 females; median age of 39.5 years, interquartile range [IQR] 30.0 to 50.3 years) and 64 COVID-19 convalescent subjects (26 males, 38 females; median age of 37 years, IQR 30.8 to 48.0). Individuals in the two cohorts were infected in the early pandemic (March 2020 to May 2021 for Post-COVID patients and February to May 2020 for the control group), when only Wuhan-like (D614G) and Alpha strain infections were present (7). Approval was obtained from the ethical committee of the Faculty for Medicine, University of Regensburg, Germany (Ref.no. 20-1785-101 and 20-1896-101). Post-COVID and control cohorts were matched regarding sex, age and time interval from symptom onset of COVID-19 to serum donation, which took place at a median of seven months post symptom onset (IQR of 5.0 to 9.0 and 6.5 to 7 months, respectively). A total of 10 out of 64 Post-COVID patients (15.6%) were vaccinated at the time of serum sampling. The majority of Post-COVID patients suffered from fatigue (88.5%), dyspnea (81.0%), and cognitive and memory impairments (71.4%).

For detection of anti-Spike antibodies of the different IgG subtypes we used a SARS-CoV-2 pre-fusion stabilized spike protein ectodomain (S ecto)-based ELISA (8). To be able to compare the different levels of IgG subtypes, we established four subtype variants of the monoclonal antibody CR3022 as standard for the respective subtype-specific ELISA (9).

We found differences in the absolute levels of the four IgG subtypes for both cohorts (**Figure 1 A**). IgG1 is by far the most abundant subtype with median concentrations of 41.4 µg/ml (IQR of 29.5 to 209.4 µg/ml) and 25.5 µg/ml (IQR of 21.9 to 32.5 µg/ml) for the Post-COVID and control cohort. These serum concentrations match well with the concentrations described by Irrgang *et al*. after one immunization with the mRNA vaccine Comirnaty (median 17.3 µg/ml with IQR of 6.9 to 21.7 µg/ml). IgG2 and IgG3 levels were similar in both cohorts (medians of 3.2 µg/ml [IQR of 2.2 to 4.9 µg/ml] and 5.5 µg/ml [IQR of 3.9 to 7.3 µg/ml] for IgG2 and 5.7 µg/ml [IQR of 2.6 to 13.7 µg/ml] and 4.0 µg/ml [IQR of 2.2 to 5.9 µg/ml] for IgG3 in the Post-COVID and control cohort, respectively). However, the levels in our study were higher than those detected by Irrgang *et al*. after one immunization with Comirnaty (medians 1.3 µg/ml and 1.0 µg/ml for IgG2 and IgG3 with IQRs of 0.9 to 2.0 µg/ml and 0.6 to 2.1 µg/ml respectively). Levels of the IgG4 subtype were lowest in both cohorts, with a median of 1.6 µg/ml (IQR of 1.2 to 2.6 µg/ml for the Post-COVID and 1.0 to 2.3 µg/ml for the control cohort respectively), whereas no IgG4 was detectable with the assay described by Irrgang *et al*.. Only the difference in IgG2 between the two cohorts reached significance at an alpha level of 0.05. IgG2 has been linked to the recognition of polysaccharide structures of bacterial cell walls as well as human endogenous glycan epitopes (2,10). It is conceivable that less efficient recognition of SARS-CoV-2 S antigen glycans could contribute to prolonged antigen circulation.

**Figure 1:**
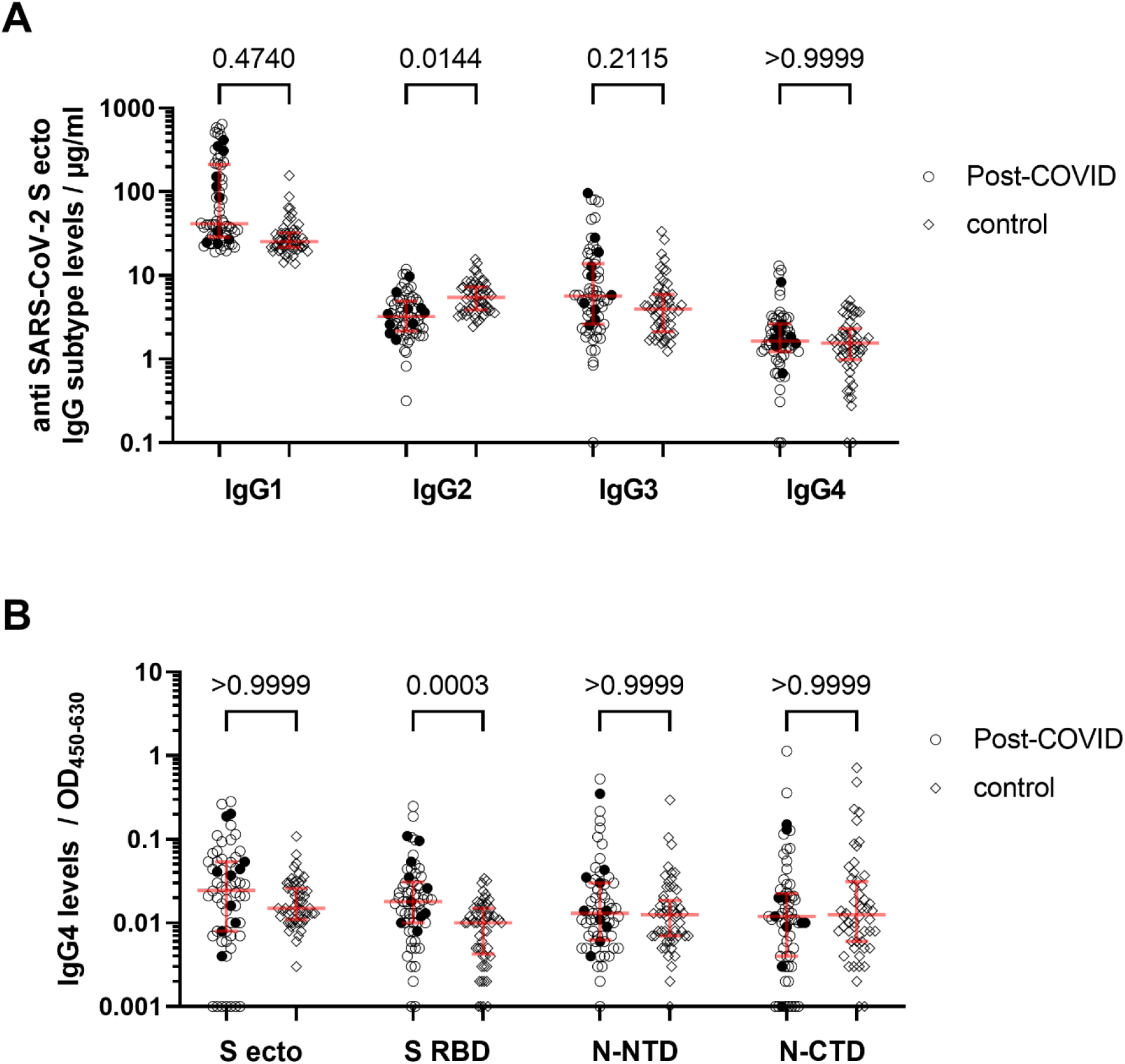
Serum IgG subtype levels in COVID-19 convalescent (n=64) and Post-COVID subjects (n=64). Serum levels of vaccinated Post-COVID patients (n=10) are indicated by filled circles. Medians and IQRs are represented in red. *P*-values of the Kruskal-Wallis test adjusted by Dunn’s *post-hoc* test are given for selected comparisons indicated by horizontal brackets. **(A)** Serum levels of IgG subtype 1-4 were quantified separately by comparison to individual subtype variants of the monoclonal antibody CR3022. The SARS-CoV-2 stabilized spike protein ectodomain (S ecto) was coated on the plate and binding of patient sera was detected by IgG subtype-specific conjugates. Serum concentration values ≤ 0 µg/ml were set to 0.1 µg/ml to allow for logarithmic scaling of the axis. **(B)** Antigen-specific IgG4 levels were determined after prolonged development of the ELISA for SARS-CoV-2’s S ecto, receptor-binding domain (RBD), and nucleocapsid N- and C-terminal domain (NTD and CTD). OD_450-600_ values ≤ 0 were set to 0.001 to allow for logarithmic scaling of the axis. For both analyses, P values changed only marginally when vaccinated Post-COVID patients were excluded.

To further investigate the impact of IgG4 antibodies in the Post-COVID context, we analyzed their reactivity against the S ecto protein, its receptor-binding domain (RBD) and the N- and C-terminal domain of the nucleocapsid-protein of SARS-CoV-2. In this assay we extended the development time of the ELISA from 4 min to 20 min to amplify the initially low signals. A small but significant difference was detected for the RBD (**Figure 1 B**), but not for the spike ectodomain and nucleocapsid proteins. Whether this difference between the cohorts reflects a pathomechanistic factor in the development of the Post-COVID syndrome is an open question. As the overall level of IgG4 is very low, we do not expect a biologically relevant role of minor differences. However, an additive effect in interaction with other factors or an impact on different subtypes of Post-COVID syndrome cannot be excluded. Furthermore, IgG4 is apparently not an appropriate biomarker for Post-COVID but may be predictive at an earlier time point, e.g. during acute infection. We were unable to answer this question because serum samples from an early post-infection time point were not available for the Post-COVID cohort.

In summary, our analysis provides insights in the relative distribution of Spike-specific IgG subtypes in Post-COVID patients and convalescent subjects not vaccinated before infection, as well as in the differences between the two cohorts and antigen-dependent differences for IgG4. Based on our dataset we do not expect IgG4 to be a central parameter in the development of Post-COVID. Rather, low IgG2 levels could contribute to the prolonged presence of SARS-CoV-2 antigens driving chronic inflammation and thus persistent Post-COVID symptoms.

## Author contributions

DP and BSc were responsible for the conception and design of the study. The Post-COVID cohort was assembled and samples generated by DS, AW, CLM, IM, AM, SB, and MP. VG, TG and DP contributed the control cohort samples. DP carried out the experimental design. JA and DP collected data. AW, CM, BSc and DP analysed and interpreted the data. Clinical symptoms of post-COVID patients were characterized by DS, TH, RB, RK, MMal, TL, and KdW with valuable intellectual input by AG, PS, MK, KR, TS, VK, BSa, MMac, and MS. The manuscript was drafted by DP and revised by BSc. All authors approved the submitted version.

## Acknowledgments

We thank all study participants for their commitment and support. We acknowledge financial support through the BMBF Project COVIDYS (project no. 01EP2105A) and the pandemic responsiveness fund of The Bavarian Ministry of Science and Art.

